# Highly sensitive mapping of *in vitro* type II topoisomerase DNA cleavage sites with SHAN-seq

**DOI:** 10.1101/2024.05.17.594727

**Authors:** Ian L. Morgan, Shannon J. McKie, Rachel Kim, Yeonee Seol, Jing Xu, Gabor Harami, Anthony Maxwell, Keir C. Neuman

## Abstract

Type II topoisomerases (topos) are a ubiquitous and essential class of enzymes that form transient enzyme-bound double-stranded breaks on DNA called cleavage complexes. The location and frequency of these cleavage complexes on DNA is important for cellular function, genomic stability, and a number of clinically important anticancer and antibacterial drugs, e.g., quinolones. We developed a simple high-accuracy end-sequencing (SHAN-seq) method to sensitively map type II topo cleavage complexes on DNA *in vitro*. Using SHAN-seq, we detected *Escherichia coli* gyrase and topoisomerase IV cleavage complexes at hundreds of sites on supercoiled pBR322 DNA, approximately one site every ten bp, with frequencies that varied by two-to-three orders of magnitude. These sites included previously identified sites and 20-50 fold more new sites. We show that the location and frequency of cleavage complexes at these sites are enzyme-specific and vary substantially in the presence of the quinolone, ciprofloxacin, but not with DNA supercoil chirality, i.e., negative vs. positive supercoiling. SHAN-seq’s exquisite sensitivity provides an unprecedented single-nucleotide resolution view of the distribution of gyrase and topoisomerase IV cleavage complexes on DNA. Moreover, the discovery that these enzymes can cleave DNA at orders of magnitude more sites than the relatively few previously known sites resolves the apparent paradox of how these enzymes resolve topological problems throughout the genome.

## Introduction

Type II topoisomerases (topos) are a ubiquitous and essential class of enzymes that form transient enzyme-bound double-stranded breaks (DSBs) on DNA called cleavage complexes^1,2^. These cleavage complexes allow the enzymes to resolve topological problems on DNA, e.g., supercoils and catenanes. However, they also pose a significant risk to genomic stability, because, if they become trapped on DNA, they can lead to DNA damage, mutations, translocations, and even cell death^3–6^. As a result, a number of small molecule inhibitors that trap cleavage complexes on DNA are clinically important anticancer and antibacterial drugs, e.g., quinolones^7–9^. Due to their wide-ranging biological and pharmacological importance, there has been considerable interest in mapping type II topo cleavage complexes on DNA and understanding the factors that influence their location and frequency^10^.

Despite the fact that topological problems need to be resolved by type II topos throughout the genome, a surprising early finding was that these enzymes tend to preferentially maintain cleavage complexes at a relatively few specific sites on DNA *in vitro* and *in vivo*^11–14^. The limitations of available methods made it difficult to precisely map the location of these sites and prevented accurate quantification of the frequency of cleavage complexes at each site. Nevertheless, analyses of cleavage site sequences revealed consensus sequences around the cleavage sites. However, in contrast to restriction enzymes, these consensus sequences were poorly defined and differed among studies. These differences were attributed to a number of factors, such as too few cleavage sites, the inclusion of “weak” cleavage sites, the use of different enzymes, and the presence and/or concentration of drugs used to trap cleavage complexes^15–18^. More broadly, the lack of well-defined consensus sequences made it difficult to accurately predict type II topo cleavage sites on DNA *de novo*. Therefore, new experimental methods were needed to accurately map the location and frequency of type II topo cleavage complexes on DNA and determine the factors that influence their site-specificity.

The advent of next-generation sequencing technologies has led to new methods for mapping the location and frequency of type II topos on DNA^10,19–25^. These methods typically fall into two categories: Chromatin Immunoprecipitation sequencing (ChIP-seq), which maps the location of type II topos bound to DNA, and end-sequencing, which maps topo-generated DSBs. Although both of these methods have led to considerable insights into the biological roles of type II topo on DNA, they have exclusively been used for *in vivo* experiments. *In vitro* experiments have several potential advantages that could help clarify the factors that influence the distribution of type II topos on DNA. First, the use of purified enzymes allows *in vitro* experiments to unambiguously determine the location and frequency of specific enzymes. Second, because cells keep cleavage levels low to maintain genomic integrity, *in vivo* experiments tend to have a low intrinsic signal-to-noise^19^. To compensate for this, *in vivo* experiments typically use drugs to trap type II topo cleavage complexes on DNA to increase their signal-to-noise. Yet, the extent to which these drugs can alter the location and frequency of type II topo cleavage complexes on DNA is unclear. In contrast, with *in vitro* experiments, high cleavage levels can be reached with or without the use of drugs. Lastly, although type II topos recognize DNA supercoiling and supercoil chirality (i.e., negative vs. positive supercoils), these factors are difficult to control and investigate *in vivo*. Recent *in vivo* studies have found that more type II topo cleavage complexes tend to be associated with positively supercoiled genomic regions than negatively supercoiled ones^19–21^. However, in *in vitro* experiments, where DNA supercoiling and supercoil chirality can be precisely controlled, type II topos tend to maintain more overall cleavage complexes on negatively supercoiled DNA than on positively supercoiled DNA^26,27^. It is unclear whether this discrepancy between *in vitro* and *in vivo* experiments is caused by a preference for genomic regions that tend to be positively supercoiled, a specific effect of DNA supercoil chirality on the enzymes’ site-specificities, or some other factor.

In this study, we developed an *in vitro* simple, high-accuracy, end-sequencing (SHAN-seq) method and used it to map *Escherichia coli (E. coli)* gyrase and topoisomerase IV (topo IV) cleavage complexes on supercoiled pBR322 plasmid DNA. For both enzymes, we detected cleavage complexes at hundreds of sites on the DNA, approximately one cleavage site every ten bp, with frequencies that varied by two-to-three orders of magnitude. We measured and compared the location and frequency of these cleavage complexes between enzymes, with and without the quinolone, ciprofloxacin (cip), and on negatively and positively supercoiled DNA. Together, our results provide an unprecedented quantitative and nucleotide-resolution view of gyrase and topo IV cleavage complexes on DNA.

## Methods

### Enzymes, DNA, and materials

*E. coli* topo IV subunits, ParC and ParE, and gyrase subunits, GyrA and GyrB, were expressed and purified as previously described^26^. The subunits were combined with 10% excess ParE over ParC or GyrB over GyrA to generate holoenzymes. All purified proteins were stored at −80°C. Negatively and positively supercoiled pBR322 DNA was purchased from Inspiralis and stored at −20°C. Analytical grade ciprofloxacin was purchased from Sigma and stored at 4°C. Proteinase K (PK), NEBNext® Ultra™ II DNA PCR-free Library Prep Kit for Illumina®, and NEBNext® Multiplex Oligos for Illumina® were purchased from NEB and stored at −20°C.

### TDP2 Purification

pET151/D-TOPO carrying N-terminal histidine tagged recombinant human TDP2 was a gift from Dr. Yves Pommier, National Cancer Institute, National Institutes of Health. The expression vector was transformed into Rosetta (DE3)pLysS competent cells (EMD Millipore). A 20 ml start-up culture containing a colony from a plate was grown overnight at 37°C and transferred to 1 L of Terrific Broth supplemented with 100 µg/ml carbenicillin and 25 µg/ml Chloramphenicol. After 3-hour incubation at 37°C, the culture was cooled to 22°C and grown overnight after addition of 1 ml of 1 M IPTG (GoldBio). Cells were harvested by centrifugation at 6000 rpm for 30 min at 4°C and resuspended in binding buffer (50 mM Tris-HCl, pH 7.5, 1% glycerol, 150 mM NaCl, 30 mM imidazole) with the addition of protease inhibitor and Benzonase (EMD Millipore). The cells were lysed by three rounds of freeze-thaw cycles followed by sonication on ice for 15 min at 30% power and high-pressure emulsification. Cell debris were removed by centrifugation at 40 krpm for 1 hour. For purification, the supernatant was applied to 5 ml of Ni-NTA agarose beads (Qiagen). The beads were washed with 200 ml of wash buffer (50 mM Tris-HCl, pH 7.5, 1% glycerol, 1 M NaCl, 30 mM imidazole, 0.01% Tween-20) and eluted with elution buffer (50 mM Tris-HCl, pH 7.5, 1% glycerol, 300 mM NaCl, 300 mM imidazole, 0.01% Tween-20). Fractions containing TDP2 were concentrated in SEC buffer (50 mM Tris-HCl, pH 7.5, 1% glycerol, 300 mM NaCl, 1 mM 2-Mercaptoethanol) using a 30 kD Amicon filter and treated with TEV protease to remove the his-tag. After collecting the removed his-tag using Ni-NTA agarose beads, the purity of TDP2 was further refined by size exclusion chromatography (Cytiva). Peak fractions of TDP2 from the size-exclusion column were concentrated in storage buffer (25 mM Tris-HCl, pH 7.5, 150 mM NaCl, 1 mM DTT, 50% glycerol) using a 30 kD Amicon filter and stored at −80°C.

### Cleavage assays and TDP2 treatment

DNA cleavage assays were carried out as previously described^26,28^. Cleavage assay mixtures contained 20 nM topo IV (heterotetramer) or 50 nM gyrase (heterotetramer) and 10 nM negatively or positively supercoiled pBR322 plasmid DNA in 50 µL of 40 mM Tris-HCl (pH 7.4), 50 mM NaCl, and 5 mM MgCl_2_ or CaCl_2_. Cip reactions contained 10 µM cip. The mixtures were incubated at 37°C for 20 min to allow cleavage complexes to form and stopped by the addition of 5 µL of 5% sodium dodecyl sulfate (SDS) and 5 µL of 250 mM Na_2_EDTA (pH 8.0). The protein was digested by adding 2 µL of PK (NEB) to the mixture and incubating at 45°C for 45 min. Protein-digested DNA was purified using a PCR Clean-up kit (Qiagen). Purified DNA was treated with 100 nM TDP2 in 30 µL of 1X rCutsmart buffer (NEB) and incubated at 37°C for 180 min to completely remove the 5’ phosphotyrosine adducts. The TDP2-treated DNA was purified using a PCR Clean-up kit (Qiagen). The TDP2-treated DNA was spiked with three synthetic DNA oligonucleotides at a molar ratio of 1:100, 2:100, and 4:100 to allow for accurate comparison of read counts across samples after sequencing.

### SHAN-seq library preparation and sequencing

The DNA was prepared for end-sequencing with two rounds of NEBNext® Ultra™ II DNA PCR-free Library Prep Kit for Illumina® and NEBNext® Multiplex Oligos for Illumina® (Unique Dual Index UMI Adapters DNA Set 1) according to the manufacturer’s instructions. Each round of library prep consisted of an end-repair step, an A-tailing step, and an adaptor ligation step. The TDP2-treated topo II-cleaved DNA was subject to a round of library prep to label the topo II-cleaved ends with sequencing adapters. The labelled DNA was fragmented to an average length of 500 bp with an ME220 focused-ultrasonicator (Covaris). The fragmented DNA was then subject to another round of library prep to label the fragmented ends with sequencing adapters. All cleanup and size selection steps were performed with AMPureXP beads (Beckman Coulter). The concentration of adapter-labelled DNA fragments in each sample was measured via qPCR using NEBNext® Library Quant Kit for Illumina® (NEB). The samples were then pooled together to an equimolar concentration and checked on a Tapestation (Agilent).

### Sequencing and data analysis of SHAN-seq libraries

Pooled SHAN-seq libraries were sequenced on an Illumina NovaSeq 6000 system (SP flow cell, 300 million reads total) with pair-end read lengths of 50bp. Reads were trimmed with Trimmomatic^29^ and mapped to the pBR322 plasmid reference sequences with BWA-MEM^30^. BAM files were sorted and indexed with SAMtools^31^. All subsequent analysis was performed with custom-written Python scripts using statistical tests from Scipy^32^. Raw plasmid read counts were extracted from BAM files using Pysam (https://github.com/pysam-developers/pysam) and normalized by spike-in DNA read counts. Cleavage sites were identified from positions with forward and reverse read counts greater than three standard deviations over background read counts. Cleavage site sequences were extracted, aligned, and used to calculate nucleotide frequencies. To account for template bias, nucleotide frequencies were compared to the overall pBR322 nucleotide frequencies. Visualizations were made with matplotlib^33^.

## Results

### Simplified High-Accuracy eNd-sequencing (SHAN-seq) method

We developed the SHAN-seq method to map type II topo cleavage complexes on DNA *in vitro* (Fig. 1A). To generate topo II cleavage complexes, we used standard *in vitro* type II topo cleavage assays^34^. In these assays, purified type II topos are incubated with DNA to form cleavage complexes, and the protein is removed from the cleavage complex using sodium dodecyl sulfate (SDS) and proteinase K (PK), revealing the topo-cleaved DNA ends. The 5’ ends of the topo-cleaved DNA are covalently attached to phosphotyrosine adducts that have been reported to block ligation to sequencing adapters^21,35^. To remove these adducts, we treated the DNA with tyrosyl DNA phosphodiesterase-2 (TDP2)^36^. After TDP2 treatment, we filled in the single-stranded overhangs and ligated the topo-cleaved DNA ends to sequencing adapters containing a unique nucleotide index. We then fragmented the DNA and ligated the fragmented ends to a second set of sequencing adapters containing a different unique nucleotide index. After pair-end sequencing, we used the unique nucleotide indices to differentiate between reads from topo-cleaved and fragmented DNA ends and selectively mapped the reads from topo-cleaved DNA ends onto the DNA reference sequence.

**Fig. 1.**
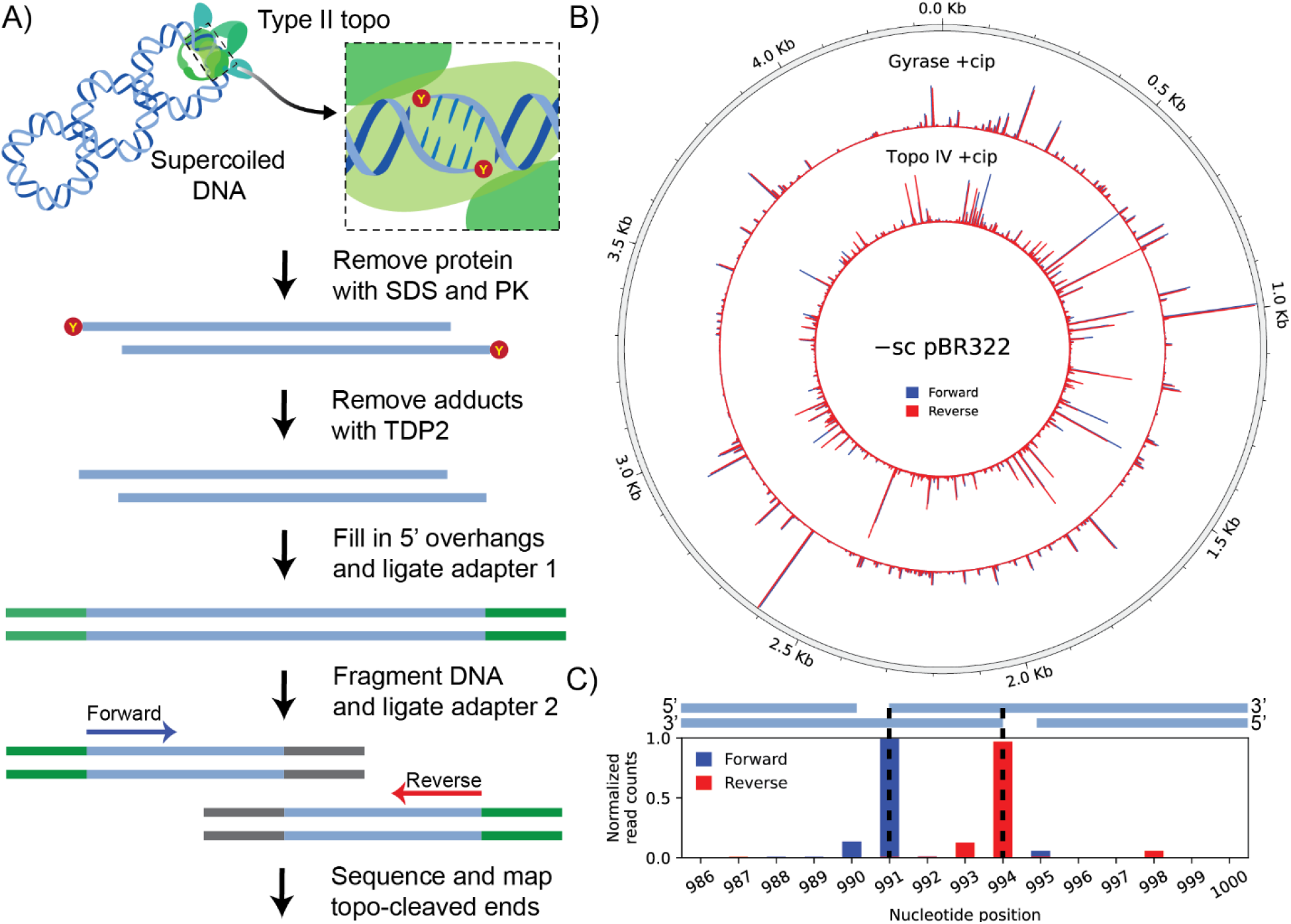
SHAN-seq sensitively maps type II topo cleavage complexes on DNA with single-nucleotide resolution. A) Schematic of SHAN-seq methodology. Adapter 1 (green) labels the topo-cleaved DNA ends and adapter 2 (grey) labels the fragmented DNA ends. (B) Cleavage maps of forward (red) and reverse (blue) read ends from *E. coli* gyrase and topo IV-cleaved negatively supercoiled (-sc) pBR322 DNA with 10 µM cip. The relative height of the bars represent the number of ends (i.e., read counts) normalized to the position with the highest read count. (C) (Upper) Cartoon of gyrase or topo IV-generated staggered DSB with four nucleotide single-stranded 5’ overhangs. (Lower) Read counts at the pBR322 strong gyrase cleavage site. Because read counts map to the first nucleotide of each read, the overhangs result in a characteristic separation between the forward and reverse read counts.

### SHAN-seq maps gyrase and topo IV cleavage complexes with single-nucleotide resolution

We tested SHAN-seq by mapping *E. coli* gyrase and topo IV cleavage complexes on negatively supercoiled pBR322 DNA. To compare our results to previous studies^13,14^, we promoted gyrase and topo IV cleavage with cip. Additionally, to maintain the topological state of the DNA, we performed our cleavage assays without ATP. Although gyrase slowly removes negative DNA supercoils in the absence of ATP^37,38^, we did not observe appreciable supercoil relaxation under our reaction conditions (Methods). After sequencing and mapping the reads from the gyrase or topo IV-cleaved DNA ends onto the plasmid reference sequence, we counted the number of read ends, i.e., read counts, at each position on the plasmid. We note that, because we did not use PCR amplification, the read counts at each position directly correspond to the number of DSBs^39^, and thus, the gyrase or topo IV cleavage levels at that position on the plasmid. For both gyrase and topo IV, the read counts from replicate libraries were highly correlated (Pearson correlation, *r* > 0.99; Supplementary Fig. 1), indicating excellent reproducibility, so they were pooled together. The resulting cleavage maps showed sharp, single-nucleotide peaks that were not present in controls lacking enzyme (Fig. 1B and Supplementary Fig. 1), indicating they were specific to the gyrase and topo IV-cleaved DNA. We observed that peaks from reads mapped in the forward and reverse orientation had a characteristic separation between them (Fig. 1C). This characteristic separation is expected because gyrase and topo IV generate staggered DSBs with four nucleotide single-stranded 5’ overhangs^10,19,21^. After shifting the reverse read counts to account for the single-stranded overhangs, the forward and reverse read counts were well-correlated (Pearson correlation, *r* > 0.9; Supplementary Fig. 1). Therefore, we identified positions that were enriched in both forward and reverse read counts as cleavage sites and counted the total read counts at each site. These results demonstrated that SHAN-seq maps the location and frequency of gyrase and topo IV cleavage complexes with single-nucleotide resolution.

### SHAN-seq sensitively detects gyrase and topo IV cleavage complexes at hundreds of sites on negatively supercoiled pBR322

In total, we identified 911 and 427 gyrase and topo IV cleavage sites on negatively supercoiled pBR322, respectively. Given that pBR322 is a 4361 bp plasmid, this means that, on average, we detected about one cleavage site every ten bp for both enzymes. The read counts across these sites varied by two-to-three orders of magnitude (Supplementary Fig. 2), suggesting that the enzymes maintain cleavage complexes at these sites with widely varying frequencies. Approximately 31% of site cleaved by gyrase were also cleaved by topo IV (Fig. 2A). However, the read counts between these two enzymes at overlapping sites were poorly correlated (*r* = 0.24, Fig. 1A), suggesting that the two enzymes have different cleavage site-specificities.

**Fig. 2.**
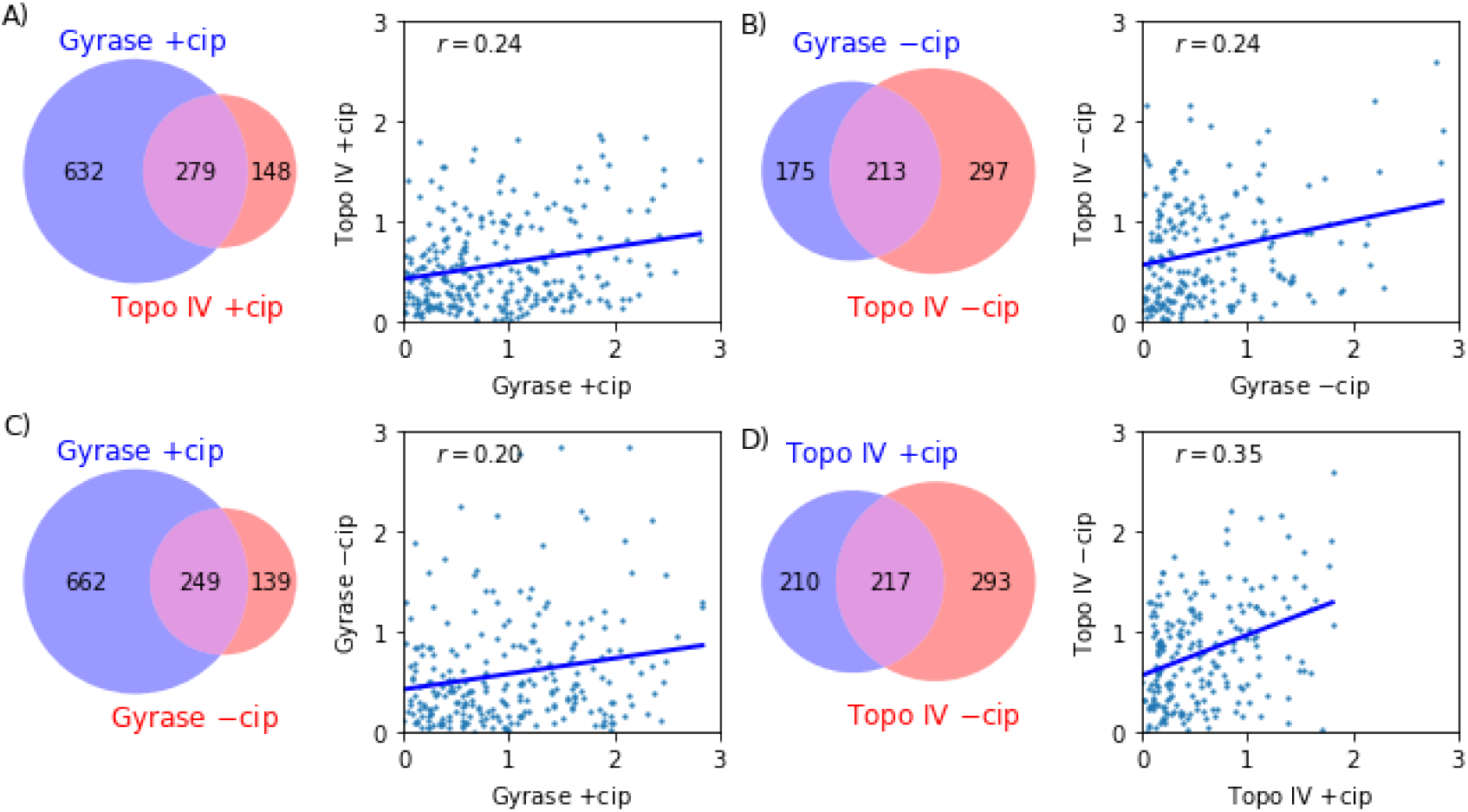
Comparisons between gyrase and topo IV cleavage sites (A) with cip and (B) without cip, and (C) gyrase and (D) topo IV with and without cip. (Left) Venn diagram of unique and shared cleavage sites. (Right) Scatterplot of log_10_ read counts for overlapping cleavage sites normalized by the site with the lowest read count. Blue lines represent best linear fit to points. Pearson correlation coefficient, *r*, for each condition is annotated.

To our knowledge, the most extensive analysis of bacterial topo II cleavage sites on pBR322 to-date used the quinolone, oxolinic acid, to identify 19 cleavage sites with nucleotide-level resolution^14^. Of these 19 sites cleavage sites, 14 and 7 of them were included among our gyrase and topo IV cleavage sites, respectively. Most of these previously identified sites had relatively high read counts. Five of the 19 sites were not included among our gyrase or topo IV cleavage sites, likely due to the indirect cloning procedure that was used to identify cleavage sites in the previous study or quinolone-specific differences^19^. Nevertheless, we captured the majority of previously identified sites, as well as an order of magnitude more sites not previously identified on pBR322^14^. These results further validate that SHAN-seq maps gyrase and topo IV cleavage complexes and indicate that it has a higher detection sensitivity than previous methods.

### Ciprofloxacin significantly alters the location and frequency of gyrase and topo IV cleavage complexes on pBR322

To investigate the effects of cip on cleavage site-specificity, we used SHAN-seq to map gyrase and topo IV cleavage complexes without cip. To increase cleavage levels without cip, we carried out calcium-promoted cleavage assays with gyrase and topo IV on negatively supercoiled pBR322. Replacing Mg^2+^ with Ca^2+^ in *in vitro* cleavage assays has been shown to increase overall cleavage levels without altering site-specificity^40^. We performed SHAN-seq on the DNA from the calcium-promoted cleavage assays and mapped the gyrase and topo IV-cleaved DNA ends onto the reference sequence. Similar to our cip results, replicate read counts were highly-correlated (Pearson correlation coefficient, r>0.99; Fig. S1), so they were pooled together, the resulting cleavage maps showed sharp, single-nucleotide peaks, and the forward and reverse read count peaks had the characteristic separation indicative of the 5’ single-stranded overhangs generated by the enzymes (Fig. S1). However, we observed that the location and frequency of the peaks on the cleavage maps without cip were distinct from the ones with cip (Fig. S1).

We detected 388 and 510 gyrase and topo IV cleavage sites, respectively, without cip on negatively supercoiled pBR322 DNA. The larger number of topo IV cleavage sites without cip likely reflects the fact that topo IV tends to have higher intrinsic cleavage levels compared to gyrase^26,27,41^. Approximately 55% of gyrase cleavage sites were also cleaved by topo IV, but the read counts at these overlapping sites were poorly correlated (Fig. 2B; Pearson correlation, *r* = 0.24). These results indicate that, even without cip, gyrase and topo IV have different cleavage site-specificities.

When we compared cleavage sites with and without cip, we found substantial differences. For gyrase, approximately 27% of cleavage sites with cip were also detected without cip (Fig. 2C), whereas, for topo IV, approximately 51% of cleavage sites with cip were also detected without cip (Fig. 2D). However, for both enzymes, the read counts at these overlapping sites were poorly correlated (Pearson correlation, *r* ≤ 0.35). These results indicate that cip substantially alters gyrase and topo IV’s cleavage site-specificities.

### DNA supercoil chirality does not affect gyrase and topo IV site-specificity

To investigate the effects of DNA supercoil chirality on cleavage site-specificity, we carried out both calcium-promoted and cip-promoted cleavage assays with gyrase and topo IV on positively supercoiled pBR322 DNA. We performed SHAN-seq on the gyrase and topo IV-cleaved DNA and mapped the topo-cleaved reads onto the plasmid reference sequence. Similar to our negatively supercoiled results, replicate libraries had read counts that were highly correlated (Pearson correlation, *r* > 0.99, Supplementary Fig. 3), so they were pooled together, the cleavage maps showed sharp, single-nucleotide peaks that were not present on controls lacking enzyme, and the forward and reverse read count peaks had the characteristic separation indicative of the single-stranded 5’ overhangs generated by the enzymes (Pearson correlation, *r* > 0.9; Supplementary Fig. 3).

In total, we detected 758 and 243 cleavages sites with cip and 345 and 208 cleavage sites with calcium on the gyrase and topo IV-cleaved positively supercoiled DNA, respectively. Similar to our results on negatively supercoiled DNA, the read counts at these sites varied by several orders of magnitude, but had lower overall read counts (Supplementary Fig. 4). The lower overall number of cleavage sites and read counts compared to our negatively supercoiled results reflects the fact that these enzymes tend to maintain fewer overall cleavage complexes on positively supercoiled DNA^26,27^. For both enzymes and conditions, most of the cleavage sites detected on positively supercoiled DNA were also detected on negatively supercoiled DNA, and the read counts at overlapping sites were highly correlated (Pearson correlation; *r* ≥ 0.75; Fig. 3). These results suggest that DNA supercoil chirality influences the overall frequency of cleavage complexes at all sites, but not the enzymes’ site-specificities, regardless of whether cleavage is promoted with cip or calcium.

**Fig. 3.**
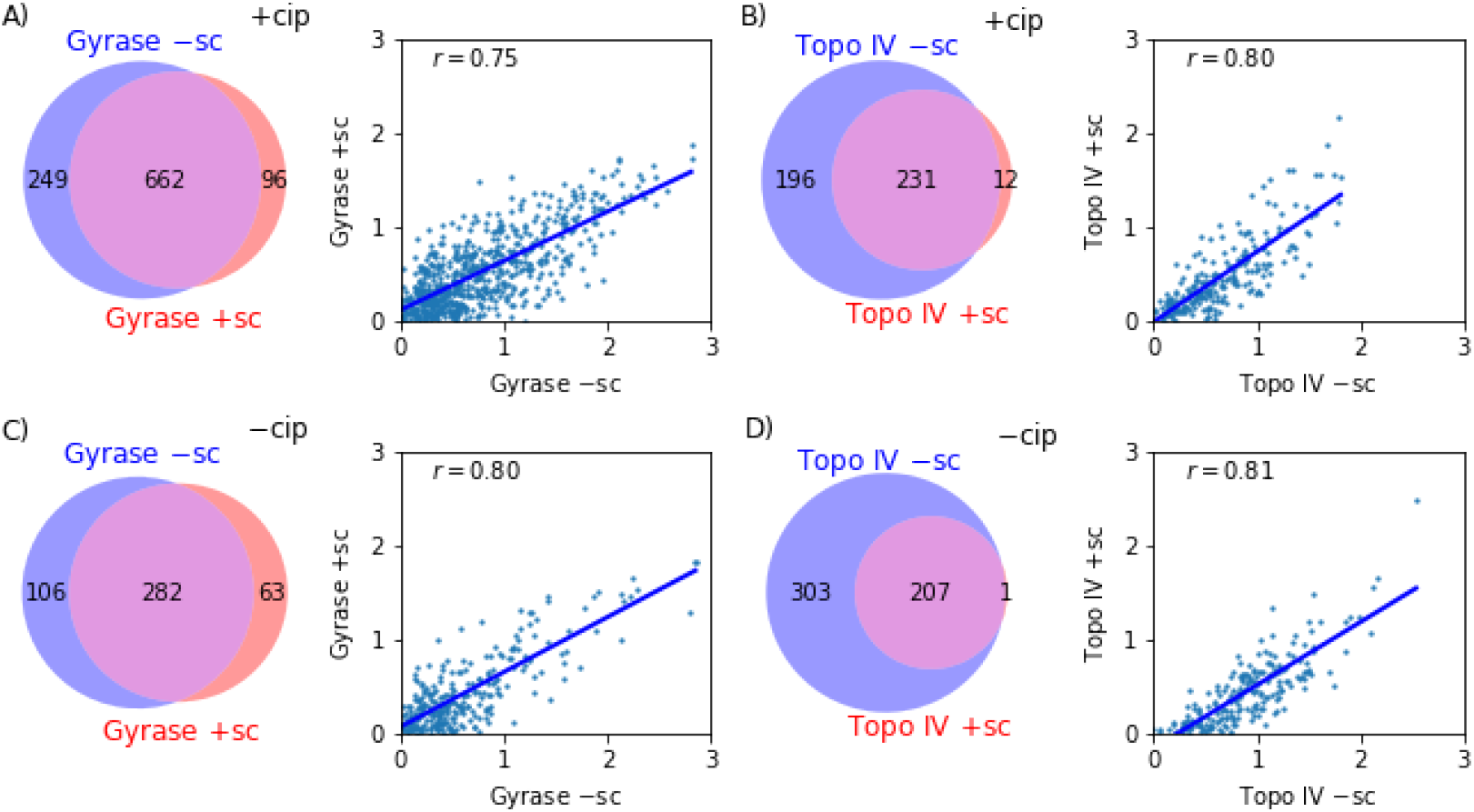
Comparison between (A and C) gyrase and (B and D) topo IV cleavage sites (A and B) with cip and (C and D) without cip between negatively supercoiled (−sc) and positively supercoiled (+sc) pBR322 DNA. (Left) Venn diagram of unique and shared cleavage sites. (Right) Scatterplot of log_10_ read counts for overlapping cleavage sites normalized by the site with the lowest read count. Blue lines represent best linear fit to points. Annotation shows Pearson correlation coefficient, *r*.

### Gyrase and topo IV exhibit similar but distinct cleavage site sequence characteristics

We used the hundreds of cleavage sites that we identified to calculate cleavage site sequence characteristics for both enzymes. We aligned the cleavage site sequences with respect to the two nicks on the top and bottom strands and calculated the frequency of each nucleotide at each position (Fig. 1C). To account for template bias, we calculated the relative difference between the nucleotide frequency at each position and the overall background nucleotide frequency of pBR322, i.e., nucleotide frequency bias. This analysis revealed a 20 nucleotide core region surrounding the cleavage site with substantial nucleotide frequency biases (Chi-squared test, *p* ≪ 0.05; Supplementary Fig. 5). Within this region, we recalculated the nucleotide frequencies to account for the inclusion of “weak sites” (i.e., low frequency sites) by weighting each cleavage site by its total read counts (Supplementary Fig. 6). The overall consensus sequences were similar for the weighted and unweighted nucleotide frequencies, so, for simplicity, we proceeded with the unweighted version (Fig. 4).

**Fig. 4.**
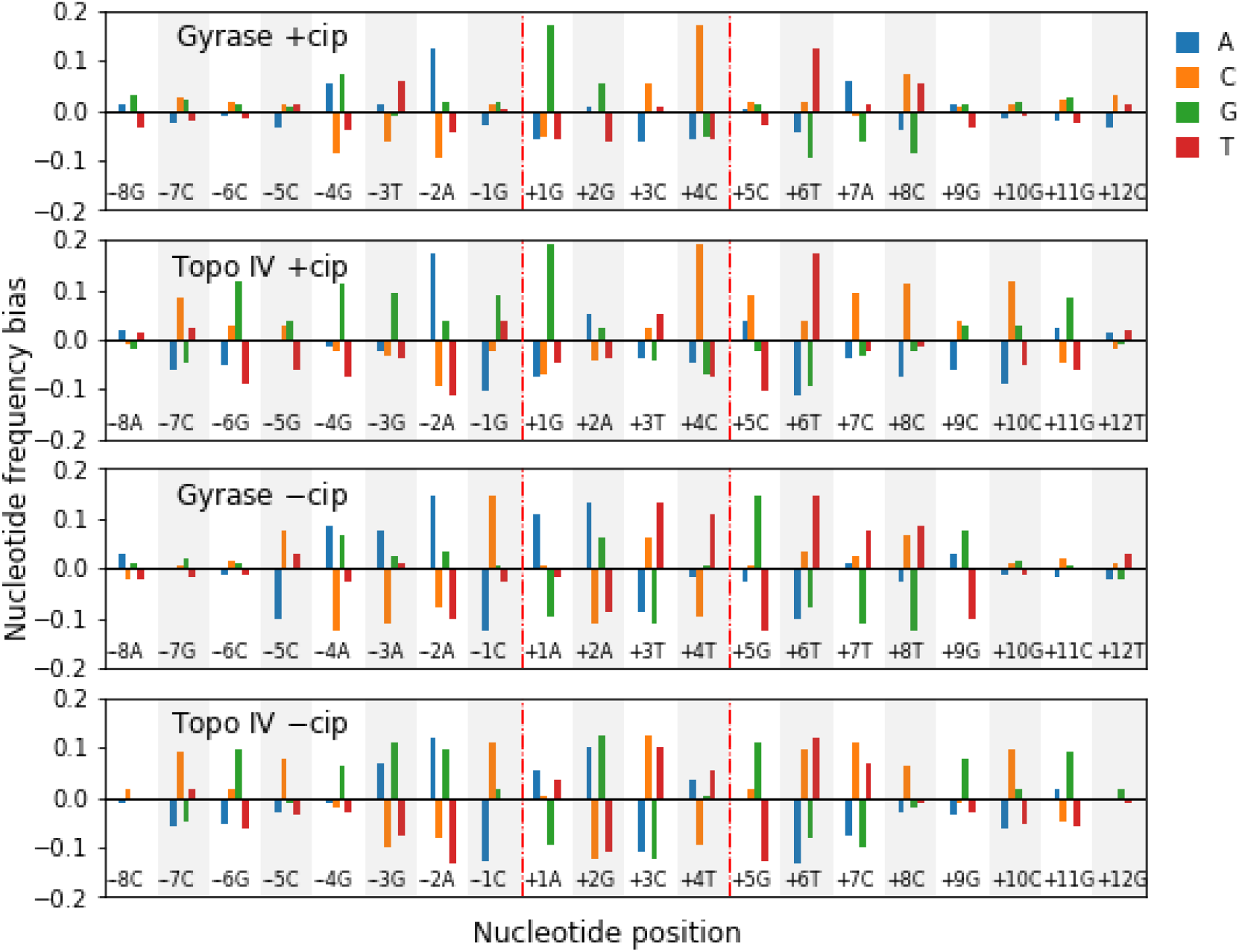
Nucleotide frequency biases for gyrase and topo IV show enzyme- and cip-specific sequence characteristics. Biases are measured as the relative change between measured and background nucleotide frequencies for each nucleotide at each position. The red vertical dashed lines denotes the position of the top and bottom strand nicks for gyrase and topo IV. Annotations represent the most frequent nucleotides at each position.

The nucleotide frequency biases within the 20 nucleotide core region were symmetric with respect to the center of the two nicks between positions +2/+3 (Fig. 4). By convention, we denoted the 5’ and 3’ nucleotide positions adjacent to the top and bottom strand nicks generated by the enzymes as −1/+1 and +4/+5. With cip, both enzymes’ cleavage sites shared a preference for −4G/+8C, −2A/+6T, and +1G/+4C. Gyrase also had preferences for −3T/+7A and +2G/+3C, whereas topo IV had preferences for −7C/+11G, −6G/+10C, −3G/+7C, and +2A/+3T. Without cip, both enzymes no longer showed a preference for +1G/+4C and had a more pronounced preference for −1C/+5G and +1A/+4T. Additionally, gyrase gained a slight preference for −4A/+8T, −3A/+7T, and +2A/+3T, whereas topo IV gained a slight preference for −5C/+9G and +2G/+3C. Overall, the largest sequence characteristic differences with and without cip tended to be adjacent to the two nicks, whereas the enzyme-specific differences were distributed throughout the core region (Supplementary Fig. 7).

In addition to the nucleotide frequency biases within the core 20 nucleotide region, we identified a weak periodic nucleotide frequency bias for gyrase, but not topo IV, flanking the core region within 60 nucleotides of the cleavage site (Supplementary Fig. 8). This periodic nucleotide bias is consistent with, albeit slightly weaker than, *in vivo* end-sequencing measurements of gyrase and topo IV cleavage sites and has been associated with the length of DNA gyrase wraps around its CTD^19,20^.

## Discussion

We have developed an *in vitro* simplified, high-accuracy end-sequencing method (SHAN-seq) and demonstrated that it can sensitively map the location and frequency of type II topo cleavage complexes on DNA. SHAN-seq’s design has several notable benefits. First, in comparison to most end-sequencing methods, it is relatively simple; it can be performed in a single day with commercially available reagents. Second, it sequences both 5’ and 3’ DNA ends. Although we used TDP2 to remove phosphotyrosine adducts from the enzyme-cleaved 5’ DNA ends, if TDP2 is not available, the reads from 3’ ends can still be used to map cleavage complexes without any modifications to the method. Lastly, it does not require PCR amplification, which can be biased^42^ and makes accurate quantification challenging^39,43^.

We used SHAN-seq to map *E. coli* gyrase and topo IV cleavage complexes on supercoiled pBR322 DNA. Surprisingly, in contrast to the relatively few sites detected in previous studies^11–14^, we detected hundreds of cleavage sites, more than one cleavage site every ten bp. These sites included previously identified sites on pBR322^14^, as well as 20-50 fold more new sites, significantly exceeding previous estimates of the density of type II topo cleavage sites on DNA^44^. At these sites, the read counts, and thus, the frequency of cleavage complexes varied by over two-to-three orders of magnitude, highlighting SHAN-seq’s exquisite detection sensitivity. The fact that most newly identified sites had relatively low cleavage levels likely accounts for why previous methods did not detect them. The wide variation in the frequency of cleavage complexes across the sites was also surprising. Because we measured the frequency of cleavage complexes at equilibrium, we cannot say whether the wide variation represents differences in binding, forward cleavage, or reverse religation rates. Ensemble biochemical studies on a limited number of cleavage sites have suggested that sites with higher cleavage levels (i.e., strong cleavage sites) tend to have higher binding affinities^19,45^. However, other *in vitro* biochemical and *in vivo* ChIP-seq experiments results conflict with these findings^46,47^. Regardless of the source of the wide variations, it is tempting to suggest that strong cleavage sites represent sites of higher enzyme activity. Yet, single-molecule studies have shown that gyrase and topo IV tend to have robust sequence-independent relaxation activity^48,49^. Moreover, in ensemble biochemical studies, introducing strong cleavage sites into a plasmid does not improve enzyme activity, and can even inhibit it^19,45^. Thus, it is likely that some, if not all, sites with lower cleavage complex frequencies can still support enzyme activity. From a biological viewpoint, the ability to form cleavage complexes at many sites could be beneficial for these enzymes’ role in regulating DNA topology throughout the genome.

Due to the low intrinsic cleavage levels of most type II topos, drugs, such as quinolones, are often used to increase and study cleavage *in vitro* and *in vivo*^11–14^. Although *in vitro* studies have indicated that these drugs can alter cleavage site sequence characteristics^15^, the full extent to which they affect the distribution of cleavage complexes on DNA was not clear. We directly compared gyrase and topo IV cleavage complexes with and without cip and found that cip substantially altered the location and relative frequency of cleavage complexes on DNA for both enzymes (Fig. 2). Notably, a ChIP-seq experiment in *E. coli* found a similarly low correlation between the frequency of topo IV bound to specific sites on DNA in the presence and absence of the quinolone, norfloxacin^47^. This finding was attributed to the difference in topo IV binding and cleavage site-specificity but could also be explained by the effects of the quinolone. Hence, although drugs are useful tools for studying type II topo cleavage complexes, results based on their use may not represent the intrinsic distribution of type II topo cleavage complexes on DNA should be interpreted with caution. Conversely, in addition to increasing cleavage levels, the extent to which these drugs alter cleavage site-specificity may be important for their pharmacological action.

In addition to the effects of quinolones, we found that gyrase and topo IV have distinct enzyme-specific cleavage site-specificities. Although previous *in vitro* and *in vivo* studies have found differences in gyrase and topo IV cleavage patterns^16,20^, they have only been compared in the presence of quinolones. Given that quinolones have differential effects on gyrase and topo IV^50^, these effects could have accounted for the apparent difference in their cleavage patterns. However, our results unambiguously show that the enzymes have different cleavage site-specificities both in the presence and absence of cip. This suggests that cleavage site-specificity is encoded into the enzymes’ sequence and structure.

Despite the fact that gyrase and, to a lesser extent, topo IV recognize DNA supercoil chirality^26,27^, maintaining fewer overall cleavage complexes on negatively supercoiled DNA, we found that DNA supercoil chirality did not affect either enzymes’ cleavage site-specificity (Fig. 3). This result agrees with a previous low resolution *in vitro* study on *S. pneumoniae* gyrase and topo IV in the presence of quinolones that found that DNA supercoiling and supercoil chirality did not qualitatively influence gyrase or topo IV cleavage patterns^16^. Although we cannot rule out that some individual cleavage sites may still respond to DNA supercoil chirality, our results indicate that it largely serves to independently and globally tune the overall frequency of gyrase and topo IV cleavage complexes on DNA for all cleavage sites. Therefore, supercoiling effects on gyrase and topo IV cleavage complex site-specificity cannot account for why *in vivo* experiments have found more cleavage complexes on genomic regions that tend to be positively supercoiled^19–21^. Instead, these findings may be due to the differential binding affinity of the enzymes on negatively and positively supercoiled DNA or the compensatory action of type I topos^51^.

We used the hundreds of cleavage sites detected with SHAN-seq to calculate consensus cleavage site sequence characteristics for gyrase and topo IV with and without cip (Fig. 4). Despite the inclusion of hundreds of sites and accounting for the template bias, the presence of drugs, and cleavage complex frequency biases, the gyrase and topo IV consensus cleavage site sequence characteristics are still relatively weak and typically allow for multiple nucleotides at each position. Nevertheless, we identified a 20 nucleotide core regions around the cleavage sites with significant nucleotide frequency biases (Supplementary Fig. 5). In agreement with numerous *in vitro* and *in vivo* studies that derived cleavage site consensus sequences in the presence of quinolones^14–17,19,20^, we found that both enzymes’ sequence characteristics had strong preferences for −4G/+8C, −2A/+6T, and +1G/+4C in the presence of cip (Fig. 4). Additionally, our vast number of cleavage sites allowed us to further refine several additional sequence characteristics within the 20 nucleotide core region for gyrase and topo IV. To our knowledge, only one study has investigated sequence characteristics for topo IV in the absence of small molecule inhibitors^15^. In agreement with this study, our results in the absence of cip showed that gyrase and topo IV maintained a strong preference for −4G/+8C and −2A/+6T and lost the strong preference for +1G/+4C. We also resolved an additional shift from −1G/+5C to −1C/+5G for both enzymes. These results indicate that cip primarily alters the preferred nucleotides adjacent to the DNA breaks. Structures of gyrase and topo IV with quinolones show two drug molecules^52,53^, one intercalated between the −1 and +1 nucleotides and another intercalated between the +4 and +5 nucleotides. Based on these structures and other biochemical evidence^54^, we attribute the shift in the sequence preferences of these nucleotides to direct interactions between the nucleotides adjacent to the breaks and the drug molecules, whereas we attribute the other apparent sequence preferences to the multiple interactions between residues within the enzyme’s active site and the DNA surrounding the cleavage site.

Notably, our novel gyrase sequence characteristics in the absence of cip was similar to previously reported sequence characteristics for gyrase with microcin B17^19^, an inhibitor with a different mechanism from quinolones. These findings suggest that inhibitors with a different mechanisms than quinolones or other means of increasing cleavage levels (e.g., hypercleavage type II topo mutants^5,6^) may be useful for determining the intrinsic distribution of type II topo cleavage complexes on DNA *in vivo*.

Despite the vast differences in the location and frequency of gyrase and topo IV cleavage complexes both with and without cipro, we were surprised to find only modest differences between the two enzymes’ sequence characteristics throughout the core 20 nucleotide region (Supplementary Fig. 7). These weak differences could be caused by subtle differences in the contacts between residues in the enzymes’ active sites and the DNA around the cleavage site. In addition, we identified weak periodic nucleotide frequency biases within 60 nucleotides of the cleavage site for gyrase but not topo IV (Supplementary Fig. 8). These periodic sequence characteristics are consistent with previous findings from *in vivo* end-sequencing experiments that have been attributed to gyrase wrapping DNA around its CTD^19,20^. The slightly weaker biases in our results compared to *in vivo* studies is likely due to the presence of chloride ions in our cleavage assays. *In vitro* biochemical assays have shown that chloride ions can partially inhibit gyrase’s ability to wrap DNA around its CTD^55^. However, given the length of the periodic nucleotide biases and modest nucleotide frequency differences within the core region, we attribute the vast difference in the location and frequency of the enzymes’ cleavage complexes primarily to gyrase’s ability to wrap DNA around its CTD. Future experiments with gyrase mutants lacking the CTD could definitively differentiate the contribution of its CTD and core region to its site-specificity.

More broadly, our results provide an unprecedented quantitative and nucleotide-resolution view of *E. coli* gyrase and topo IV cleavage complexes on DNA and demonstrate the power and versatility of SHAN-seq as a highly sensitive *in vitro* end-sequencing method. The same methodology is applicable to other bacterial and eukaryotic type II topos, type I topos, and the vast array of other site-specific nucleases and recombinases. For many of these enzymes, their cleavage site-specificity and/or the effects of drugs and DNA topology is largely unknown but likely to be important. For example, recent evidence has indicated that DNA supercoiling alters the cleavage site-specificity of the Cas9 nuclease^56^. Therefore, we expect that SHAN-seq will be a useful and broadly applicable tool for mapping DNA cleavage *in vitro* that complements existing *in vivo* methods.

## Acknowledgements

We thank James Berger for providing the *E. coli* gyrase plasmid and Yves Pommier for providing the TDP2 plasmid. We thank Bobby Hogg, Joseph Chapman, and the members of the Neuman lab for helpful discussions. This work was funded by the intramural research program of the National Heart, Lung, and Blood Institute of the National Institutes of Health (1ZIAHL001056). I.L.M. was supported by a National Institute of General Medical Sciences fellowship (1FI2GM142536). S.J.M was supported by a Wellcome Trust fellowship (200928/Z16/Z). A.M. was supported by Biotechnology and Biosciences Research Council (UK) Institute Strategic Programme (BB/P012523/1) and Wellcome Trust (110072/Z/15/Z) awards. This work utilized the resources of the NHLBI DNA Sequencing and Genomics Core and NIH HPC Biowulf cluster (http://hpc.nih.gov).

## Data and code availability

The data and code for all figures are deposited on the Figshare database (10.25444/nhlbi.25789329). High-throughput sequencing data are available from the NCBI GEO database (GSE267149).

## Author Contributions

I.L.M., S.J.M, and K.C.N conceived the project and developed the SHAN-seq methodology. I.L.M and R.K. generated SHAN-seq libraries. Y.S. and G.H. expressed and purified proteins. I.L.M. and J.X. developed the analysis pipeline. I.L.M interpreted the observations and wrote the manuscript. K.C.N., Y.S., S.J.M, and A.M. reviewed and edited the manuscript.

## References

1. McKie, S. J., Neuman, K. C. & Maxwell, A. DNA topoisomerases: Advances in understanding of cellular roles and multi-protein complexes via structure-function analysis. BioEssays 43, 2000286 (2021).

2. Bush, N. G., Evans-Roberts, K. & Maxwell, A. DNA Topoisomerases. EcoSal Plus 6, ecosalplus.ESP-0010-2014 (2015).

3. Nitiss, J. L. DNA Topoisomerases in DNA Repair and DNA Damage Tolerance. in DNA Damage and Repair (eds. Nickoloff, J. A. & Hoekstra, M. F. 517–537 (Humana Press, Totowa, NJ, 1998). doi:10.1007/978-1-59259-455-9_23.

4. Pommier, Y., Nussenzweig, A., Takeda, S. & Austin, C. Human topoisomerases and their roles in genome stability and organization. Nat Rev Mol Cell Biol (2022) doi:10.1038/s41580-022-00452-3.

5. Stantial, N. et al. Trapped topoisomerase II initiates formation of de novo duplications via the nonhomologous end-joining pathway in yeast. Proc. Natl. Acad. Sci. U.S.A. 117, 26876–26884 (2020).

6. Boot, A. et al. Recurrent mutations in topoisomerase IIα cause a previously undescribed mutator phenotype in human cancers. Proc. Natl. Acad. Sci. U.S.A. 119, e2114024119 (2022).

7. Bush, N. G., Diez-Santos, I., Abbot, L. R. & Maxwell, A. Quinolones: Mechanism, Lethality and Their Contributions to Antibiotic Resistance. Molecules 25, 5662 (2020).

8. Aldred, K. J., Kerns, R. J. & Osheroff, N. Mechanism of Quinolone Action and Resistance. Biochemistry 53, 1565–1574 (2014).

9. Pommier, Y. Drugging Topoisomerases: Lessons and Challenges. ACS Chem. Biol. 8, 82–95 (2013).

10. McKie, S. J., Maxwell, A. & Neuman, K. C. Mapping DNA Topoisomerase Binding and Cleavage Genome Wide Using Next-Generation Sequencing Techniques. Genes 11, 92 (2020).

11. Kirkegaard, K. & Wang, J. C. Mapping the topography of DNA wrapped around gyrase by nucleolytic and chemical probing of complexes of unique DNA sequences. Cell 23, 721–729 (1981).

12. Fisher, L. M., Mizuuchi, K., O’Dea, M. H., Ohmori, H. & Gellert, M. Site-specific interaction of DNA gyrase with DNA. Proc. Natl. Acad. Sci. U.S.A. 78, 4165–4169 (1981).

13. Morrison, A. & Cozzarelli, N. R. Site-specific cleavage of DNA by E. coli DNA gyrase. Cell 17, 175–184 (1979).

14. Lockshon, D. & Morris, D. R. Sites of reaction of Escherichia coli DNA gyrase on pBR322 in vivo as revealed by oxolinic acid-induced plasmid linearization. Journal of Molecular Biology 181, 63–74 (1985).

15. Arnoldi, E.Pan, X.-S. & Fisher, L. M. Functional determinants of gate-DNA selection and cleavage by bacterial type II topoisomerases. Nucleic Acids Research 41, 9411–9423 (2013).

16. Leo, E. et al. Novel Symmetric and Asymmetric DNA Scission Determinants for Streptococcus pneumoniae Topoisomerase IV and Gyrase Are Clustered at the DNA Breakage Site. Journal of Biological Chemistry 280, 14252–14263 (2005).

17. Richter, S. N. et al. Hot-spot consensus of fluoroquinolone-mediated DNA cleavage by Gram-negative and Gram-positive type II DNA topoisomerases. Nucleic Acids Research 35, 6075–6085 (2007).

18. Capranico, G. & Binaschi, M. DNA sequence selectivity of topoisomerases and topoisomerase poisons. Biochimica et Biophysica Acta (BBA) - Gene Structure and Expression 1400, 185–194 (1998).

19. Sutormin, D., Rubanova, N., Logacheva, M., Ghilarov, D. & Severinov, K. Single-nucleotide-resolution mapping of DNA gyrase cleavage sites across the Escherichia coli genome. Nucleic Acids Research 47, 1373–1388 (2019).

20. Sutormin, D., Galivondzhyan, A., Gafurov, A. & Severinov, K. Single-nucleotide resolution detection of Topo IV cleavage activity in the Escherichia coli genome with Topo-Seq. Front. Microbiol. 14, 1160736 (2023).

21. Gitens, W. H. et al. A nucleotide resolution map of Top2-linked DNA breaks in the yeast and human genome. Nat Commun 10, 4846 (2019).

22. Canela, A. et al. DNA Breaks and End Resection Measured Genome-wide by End Sequencing. Molecular Cell 63, 898–911 (2016).

23. Baranello, L. et al. DNA Break Mapping Reveals Topoisomerase II Activity Genome-Wide. IJMS 15, 13111–13122 (2014).

24. Miller, E. L. et al. TOP2 synergizes with BAF chromatin remodeling for both resolution and formation of facultative heterochromatin. Nat Struct Mol Biol 24, 344–352 (2017).

25. Cowell, I. G. et al. Model for MLL translocations in therapy-related leukemia involving topoisomerase IIβ-mediated DNA strand breaks and gene proximity. Proc. Natl. Acad. Sci. U.S.A. 109, 8989–8994 (2012).

26. Ashley, R. E. et al. Activities of gyrase and topoisomerase IV on positively supercoiled DNA. Nucleic Acids Research 45, 9611–9624 (2017).

27. Jian, J. Y. et al. Basis for the discrimination of supercoil handedness during DNA cleavage by human and bacterial type II topoisomerases. Nucleic Acids Research gkad190 (2023) doi:10.1093/nar/gkad190.

28. Aldred, K. J. et al. Role of the Water–Metal Ion Bridge in Mediating Interactions between Quinolones and Escherichia coli Topoisomerase IV. Biochemistry 53, 5558–5567 (2014).

29. Bolger, A. M., Lohse, M. & Usadel, B. Trimmomatic: a flexible trimmer for Illumina sequence data. Bioinformatics 30, 2114–2120 (2014).

30. Li, H. & Durbin, R. Fast and accurate long-read alignment with Burrows–Wheeler transform. Bioinformatics 26, 589–595 (2010).

31. Li, H. et al. The Sequence Alignment/Map format and SAMtools. Bioinformatics 25, 2078–2079 (2009).

32. Virtanen, P. et al. SciPy 1.0: fundamental algorithms for scientific computing in Python. Nat Methods 17, 261–272 (2020).

33. Hunter, J. D. Matplotlib: A 2D graphics environment. Computing in Science & Engineering 9, 90–95 (2007).

34. Burden, D. A., Froelich-Ammon, S. J. & Osheroff, N. Topoisomerase II-Mediated Cleavage of Plasmid DNA. in DNA Topoisomerase Protocols (eds. Osheroff, N. & Bjornsti, M.-A.) 283–289 (Humana Press, Totowa, NJ, 2001). doi:10.1385/1-59259-057-8:283.

35. Canela, A. et al. Topoisomerase II-Induced Chromosome Breakage and Translocation Is Determined by Chromosome Architecture and Transcriptional Activity. Molecular Cell 75, 252-266.e8 (2019).

36. Gómez-Herreros, F. et al. TDP2 protects transcription from abortive topoisomerase activity and is required for normal neural function. Nat Genet 46, 516–521 (2014).

37. Gellert, M., Mizuuchi, K., O’Dea, M. H., Itoh, T. & Tomizawa, J.-I. Nalidixic acid resistance: A second genetic character involved in DNA gyrase activity. Proceedings of the National Academy of Sciences 74, 4772–4776 (1977).

38. Sugino, A., Peebles, C. L., Kreuzer, K. N. & Cozzarelli, N. R. Mechanism of action of nalidixic acid: Purification of Escherichia coli nalA gene product and its relationship to DNA gyrase and a novel nicking-closing enzyme. Proceedings of the National Academy of Sciences 74, 4767–4771 (1977).

39. Dobbs, F. M. et al. Precision digital mapping of endogenous and induced genomic DNA breaks by INDUCE-seq. Nat Commun 13, 3989 (2022).

40. Pits, S. L. et al. Use of divalent metal ions in the DNA cleavage reaction of topoisomerase IV. Nucleic Acids Research 39, 4808–4817 (2011).

41. Khodursky, A. B., Zechiedrich, E. L. & Cozzarelli, N. R. Topoisomerase IV is a target of quinolones in Escherichia coli. Proceedings of the National Academy of Sciences 92, 11801–11805 (1995).

42. Kozarewa, I. et al. Amplification-free Illumina sequencing-library preparation facilitates improved mapping and assembly of (G+C)-biased genomes. Nat Methods 6, 291–295 (2009).

43. Zhu, Y. et al. qDSB-Seq is a general method for genome-wide quantification of DNA double-strand breaks using sequencing. Nat Commun 10, 2313 (2019).

44. Cozzarelli, N. R. DNA Gyrase and the Supercoiling of DNA. Science 207, 953–960 (1980).

45. Oram, M., Howells, A. J., Maxwell, A. & Pato, M. L. A biochemical analysis of the interaction of DNA gyrase with the bacteriophage Mu, pSC101 and pBR322 strong gyrase sites: the role of DNA sequence in modulating gyrase supercoiling and biological activity: A biochemical analysis of strong DNA gyrase sites. Molecular Microbiology 50, 333–347 (2003).

46. Mueller-Planitz, F. & Herschlag, D. DNA topoisomerase II selects DNA cleavage sites based on reactivity rather than binding affinity. Nucleic Acids Research 35, 3764–3773 (2007).

47. El Sayyed, H. et al. Mapping Topoisomerase IV Binding and Activity Sites on the E. coli Genome. PLOS Genetics 12, e1006025 (2016).

48. Neuman, K. C., Charvin, G., Bensimon, D. & Croquete, V. Mechanisms of chiral discrimination by topoisomerase IV. PNAS 106, 6986–6991 (2009).

49. Stone, M. D. et al. Chirality sensing by Escherichia coli topoisomerase IV and the mechanism of type II topoisomerases. Proceedings of the National Academy of Sciences 100, 8654–8659 (2003).

50. Blanche, F. et al. Differential behaviors of Staphylococcus aureus and Escherichia coli type II DNA topoisomerases. Antimicrob Agents Chemother 40, 2714–2720 (1996).

51. Sutormin, D. et al. Interaction between transcribing RNA polymerase and topoisomerase I prevents R-loop formation in E. coli. Nat Commun 13, 4524 (2022).

52. Laponogov, I. et al. Structural insight into the quinolone–DNA cleavage complex of type IIA topoisomerases. Nat Struct Mol Biol 16, 667–669 (2009).

53. Bax, B. D. et al. Type IIA topoisomerase inhibition by a new class of antibacterial agents. Nature 466, 935–940 (2010).

54. Bromberg, K. D., Burgin, A. B. & Osheroff, N. Quinolone Action against Human Topoisomerase IIα: Stimulation of Enzyme-Mediated Double-Stranded DNA Cleavage. Biochemistry 42, 3393–3398 (2003).

55. Hobson, M. J., Bryant, Z. & Berger, J. M. Modulated control of DNA supercoiling balance by the DNA-wrapping domain of bacterial gyrase. Nucleic Acids Research 48, 2035–2049 (2020).

56. Newton, M. D. et al. Negative DNA supercoiling induces genome-wide Cas9 off-target activity. Molecular Cell 83, 3533-3545.e5 (2023).

