## Supplementary Information for "Highly sensitive mapping of *in vitro* type II topoisomerase DNA cleavage sites with SHAN-seq"

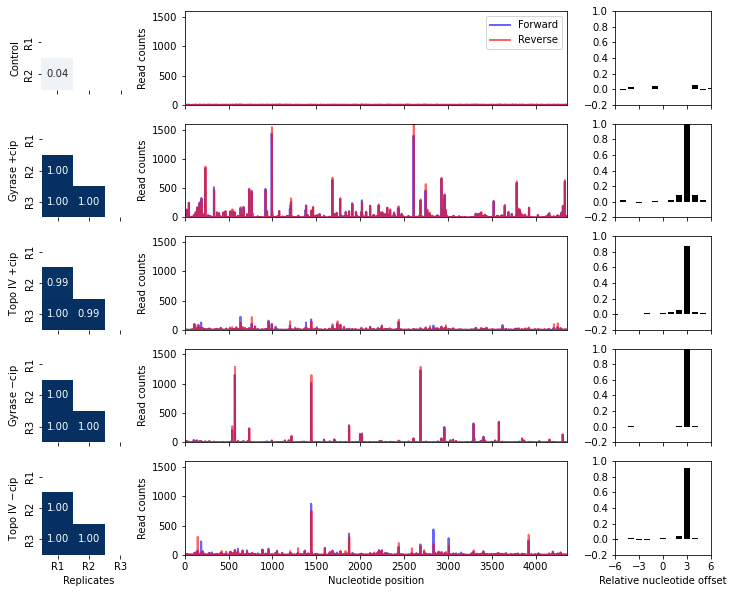

**Supplementary Figure 1:** SHAN-seq captures gyrase and topo IV cleavage sites on negatively supercoiled pBR322 DNA. (Left) Pearson correlation coefficients between replicates . (Middle) Cleavage maps of forward and reverse read counts at each nucleotide position normalized by their median value. (Right) Pearson correlation coefficient between forward and reverse read counts as a function of nucleotide offset.

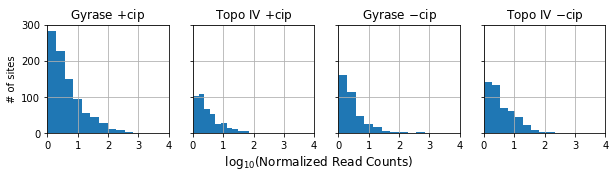

**Supplementary Figure 2:** Relative $\log_{10}$ read counts across gyrase and topo IV cleavage sites on negatively supercoiled pBR322 DNA.

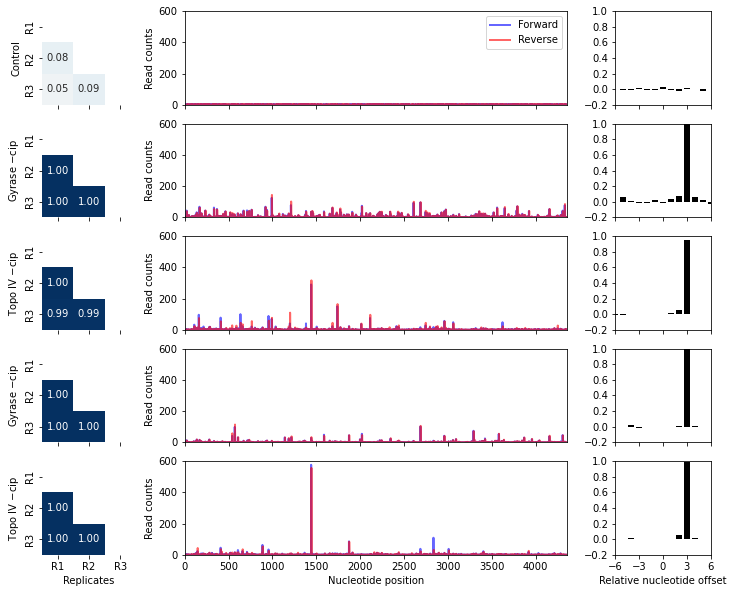

**Supplementary Figure 3:** SHAN-seq captures gyrase and topo IV cleavage sites on positively supercoiled pBR322 DNA. (Left) Pearson correlation coefficients between replicates . (Middle) Cleavage maps of normalized forward and reverse read counts at each nucleotide position normalized by their median value. (Right) Pearson correlation coefficient between forward and reverse read counts as function of nucleotide offset.

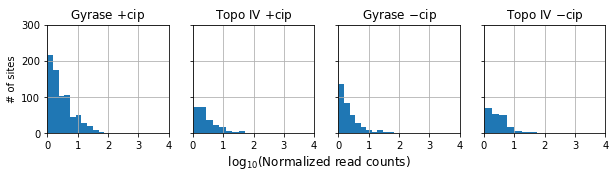

**Supplementary Figure 4:** Relative $\log_{10}$ read counts across gyrase and topo IV cleavage sites on positively supercoiled pBR322 DNA.

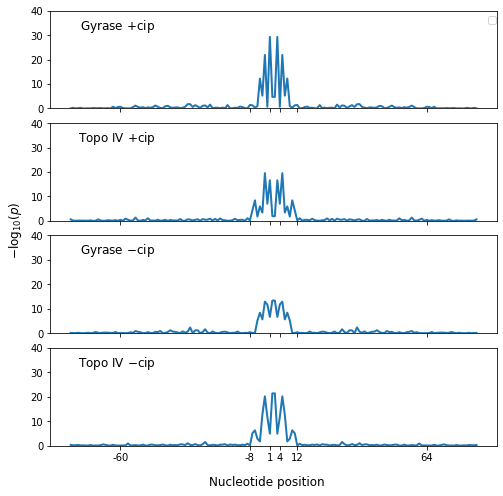

**Supplementary Figure 5: Gyrase and topo IV cleavage sites show specific sequence preferences within 20 nucleotides surrounding the cleavage site.** Values represent the negative log likelihood ($-{log}_{10}( p)$, Chi-squared test) of observing nucleotide frequency biases by chance.

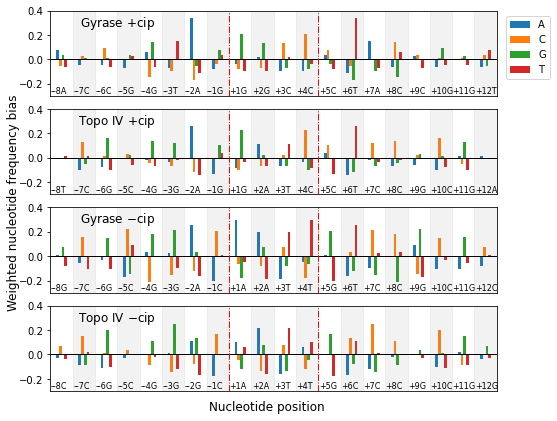

**Supplementary Figure 6: Nucleotide frequency biases weighted by read counts for gyrase and topo IV show enzyme- and cip-specific sequence characteristics.** Biases are measured as the relative change between measured and background nucleotide frequencies for each nucleotide at each position. The red vertical dashed lines denotes the position of the top and bottom strand nicks generated by gyrase and topo IV. Annotations represent the most nucleotides with the largest bias at each position.

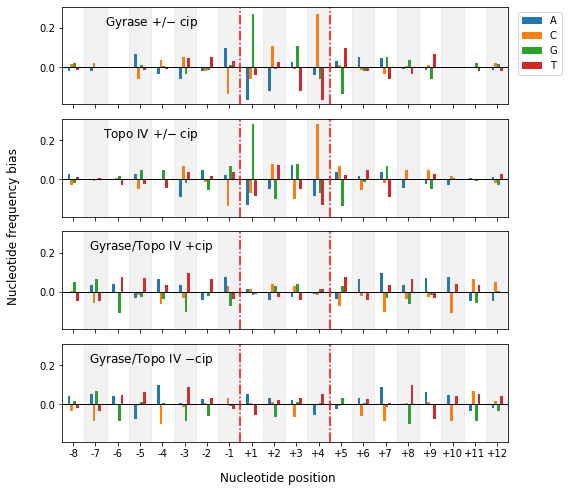

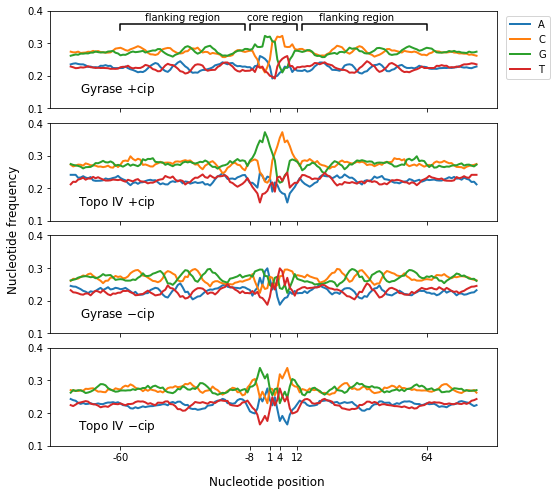

**Supplementary Figure 7: Cip and enzyme-specific nucleotide frequency differences.** Differences in nucleotide frequencies between (top) gyrase with and without cip, (upper middle) topo IV with and without cip, (lower middle) gyrase and topo IV with cip, and (bottom) gyrase and topo IV without cip.

**Supplementary Figure 8: Extended nucleotide frequencies around gyrase and topo IV cleavage sites show a periodic nucleotide frequency bias for gyrase but not topo IV.** Values indicate a five-nucleotide rolling mean of individual nucleotide frequencies from gyrase and topo IV cleavage site sequences on negatively supercoiled pBR322 DNA. The 20 nucleotide core region with strong nucleotide frequency biases and flanking regions with weaker periodic nucleotide biases for gyrase but not topo IV are marked. The pBR322 plasmid sequence has a slight overall GC bias (A/T = 0.23 and G/C = 0.27), resulting in higher cleavage site GC nucleotide frequencies.
